# Multiple indole glucosinolates and myrosinases defend Arabidopsis against *Tetranychus urticae* herbivory

**DOI:** 10.1101/2021.02.03.429630

**Authors:** Emilie Widemann, Kristie Bruinsma, Brendan Walshe-Roussel, Repon Kumer Saha, David Letwin, Vladimir Zhurov, Mark A. Bernards, Miodrag Grbić, Vojislava Grbić

**Affiliations:** Department of Biology, The University of Western Ontario, 1151 Richmond Street, London, ON N6A 5B7, Canada; Department of Microbiology and Immunology, Schulich School of Medicine and Dentistry, The University of Western Ontario, London, ON N6A 3K7, Canada; Natural and Non-Prescription Health Products Directorate Health Canada, 250 Lanark Ave, Ottawa, ON, K1A 0K9, Canada

**Keywords:** two-spotted spider mite, chemoprotection, herbivory, defenses, jasmonates, feeding suppressants

## Abstract

Arabidopsis defenses against herbivores are regulated by the jasmonate hormonal signaling pathway, which leads to the production of a plethora of defense compounds, including tryptophan-derived metabolites produced through CYP79B2/CYP79B3. Jasmonate signaling and CYP79B2/CYP79B3 limit Arabidopsis infestation by the generalist herbivore two-spotted spider mite, *Tetranychus urticae*. However, the phytochemicals responsible for Arabidopsis protection against *T. urticae* are unknown. Here, using Arabidopsis mutants that disrupt metabolic pathways downstream of CYP79B2/CYP79B3, and synthetic indole glucosinolates, we identified phytochemicals involved in the defense against *T. urticae*. We show that Trp-derived metabolites depending on CYP71A12 and CYP71A13 are not affecting mite herbivory. Instead, the supplementation of *cyp79b2 cyp79b3* mutant leaves with the 3-indolylmethyl glucosinolate and its derived metabolites demonstrated that the indole glucosinolate pathway is sufficient to assure CYP79B2/CYP79B3-mediated defenses against *T. urticae*. We demonstrate that three indole glucosinolates can limit *T. urticae* herbivory, but that they have to be processed by the myrosinases to hinder *T. urticae* oviposition. Finally, the supplementation of the mutant *myc2 myc3 myc4* with indole glucosinolates indicated that the transcription factors MYC2/MYC3/MYC4 induce additional indole glucosinolate-independent defenses that control *T. urticae* herbivory. Together, these results reveal the complexity of Arabidopsis defenses against *T. urticae* that rely on multiple indole glucosinolates, specific myrosinases, and additional MYC2/MYC3/MYC4-dependent defenses.

**One sentence summary:** Three indole glucosinolates and the myrosinases TGG1/TGG2 help protect *Arabidopsis thaliana* against the herbivory of the two-spotted spider mite *Tetranychus urticae*.

## INTRODUCTION

The jasmonate (JA) hormonal pathway is a conserved inducer of anti-herbivory defenses in a wide range of plants. In response to herbivory, JA accumulates and triggers JA-induced defense responses that vary in different plant species and include the synthesis of defensive metabolites, volatiles, and/or proteins (Howe and Jander 2008; Lortzing and Steppuhn 2016; Wasternack and Strnad 2018; Wang et al., 2019). In *Arabidopsis thaliana*, JA signaling is mediated by the MYC2, MYC3, and MYC4 transcription factors that activate a wide range of defense-associated genes (Schweizer et al., 2013). Glucosinolates are defense compounds found primarily in Brassicaceae species (Fahey et al., 2001), including Arabidopsis, whose synthesis is regulated by direct interaction between MYC2/MYC3/MYC4 and MYB transcription factors (Schweizer et al., 2013). Glucosinolates are synthesized from amino acids. Aliphatic and indole glucosinolates, derived from methionine and tryptophan respectively, are the most abundant glucosinolates in Arabidopsis (Brown et al., 2003). They are represented by a family of related compounds with 13 aliphatic and 3 stable indole metabolites (Brown et al., 2003; Mahmut 2020). These compounds have varying defensive specificities, so that some herbivores, like *Manduca sexta* and *Trichoplusia ni* are sensitive to aliphatic glucosinolates (Müller et al., 2010), some like *Myzus persicae* to indole glucosinolates (Kim and Jander 2007) and some, like *Spodoptera littoralis* and *Mamestra brassicae*, to both classes of compounds (Müller et al., 2010; Jeschke et al., 2017). Synthesized glucosinolates have to undergo further modifications to become biologically active. They are hydrolyzed by beta-glucosidases referred to as myrosinases (Bjorkman 1976; Bhat and Vyas 2019). Classical myrosinases TGG1 and TGG2 cleave the glucose group from a glucosinolate and release a highly reactive aglycone that gives rise to isothiocyanates, nitriles, epithionitriles, or thiocyanates (Barth and Jander 2006; Bones and Rossiter 2006; Blažević et al., 2020). Glucosinolates and the myrosinases TGG1 and TGG2 accumulate at high levels in different cell types, so tissue damage that is associated with herbivory is required for their contact (Xue et al., 1995; Husebye et al., 2002; Barth and Jander 2006; Ueda et al., 2006; Zhao et al., 2008; Sønderby et al., 2010; Shroff et al., 2015). Besides classical myrosinases, additional beta-glucosidases (BGLU) were demonstrated to have myrosinase activities against indole glucosinolates (Bednarek et al., 2009; Nakano et al., 2017; Nakazaki et al., 2019). They co-localize with glucosinolates but accumulate in different cell compartments.

The chelicerate *Tetranychus urticae* (the two-spotted spider mite) is an extreme generalist herbivore that uses its stylet to transverse the leaf epidermis and reach leaf mesophyll where it feeds from individual cells (Bensoussan et al., 2016). Mite feeding on Arabidopsis triggers the accumulation of JA and the induction of MYC2/MYC3/MYC4-mediated responses (Zhurov et al., 2014). Aliphatic glucosinolates are not effective against mites, however, mite fitness increases on the *cyp79b2 cyp79b3* mutant plants, indicating that the Trp-derived secondary metabolite(s) restrict mite herbivory (Zhurov et al., 2014). CYP79B2 and CYP79B3 are required for the conversion of tryptophan to indole-3-acetaldoxime (IAOx) that is further processed by CYP71A13, CYP71A12, and CYP83B1 to initiate biosynthesis of camalexin, cyanogenic metabolite 4-OH-ICN and indole glucosinolate (IG) defense compounds, respectively (Zhao et al., 2002; Sanchez-Vallet et al., 2010; Rajniak et al., 2015; Vik et al., 2018; Glindemann et al., 2019; Pastorczyk et al., 2020). 3-Indolylmethyl glucosinolate (I3M) is a parental indole glucosinolate that is further hydroxylated by cytochromes P450 of the CYP81 family and methylated by IG methyltransferases. If modifications occur on the nitrogen of the indole ring, I3M gives rise to 1OH-I3M (intermediate that does not accumulate in Arabidopsis) and 1-Methoxy-I3M (1MO-I3M). If carbon 4 of the indole ring is modified, 4OH-I3M and 4MO-I3M are synthesized (Rask et al., 2000; Meier et al., 2019).

While it was established that the JA pathway and CYP79B2/CYP79B3 are required for Arabidopsis defenses against *T. urticae*, the identity of the Trp-derived metabolite(s) remained elusive. The mutant *pad3*, deficient in the last step of camalexin production, is not more sensitive to mite infestation than wild-type (WT) plants (Zhurov et al., 2014). However, the existence of other potential defensive compounds against mites derived from Trp via CYP71A13- or CYP71A12-dependent pathways has not been tested. Furthermore, even though it was demonstrated that I3M and 1MO-I3M accumulate in mite-infested Arabidopsis leaves (Zhurov et al., 2014), their effects on mite fitness have not been demonstrated. In this study, we used a collection of Arabidopsis mutants to identify which of the Trp-derived pathways protect Arabidopsis against mite herbivory. We demonstrate that of the three CYP79B2/CYP79B3-dependent pathways, the I3M, 1MO-I3M, and 4MO-I3M glucosinolate metabolites are sufficient to protect Arabidopsis plants against *T. urticae*. We show that intact glucosinolates cannot limit mite’s ability to feed on Arabidopsis, but that they require further modifications with TGG1/TGG2 myrosinases and other currently unknown Arabidopsis factors to gain defensive activity. Furthermore, we demonstrate that in addition to Trp-derived defensive metabolites, Arabidopsis synthesizes additional indole glucosinolate-independent defenses. Our work establishes the complexity of Arabidopsis defenses that shape the interaction between Arabidopsis and generalist *T. urticae*.

## RESULTS

### Trp-derived metabolites suppress feeding of adult mites

To determine the effect of Trp-derived metabolites on mites, we performed mite feeding experiments on *cyp79b2 cyp79b3* leaves that lack Trp-derived metabolites and fully defended Columbia-0 (Col-0) wild-type leaves. Repellent activity of Trp-derived defenses was challenged using a choice experiment where adult female mites could have selected either a *cyp79b2 cyp79b3* or a Col-0 leaf to feed on. To track leaf genotypes mites fed on, we stained one of the two leaves with blue dye. When mites fed on blue-stained leaves they excreted blue feces, Fig. 1A, so the frequency of blue *vs*. normal feces then allowed us to distinguish and quantify mite feeding on individual leaves. A similar number of blue and non-stained feces in control experiments with leaves of the same genotype indicated that the blue staining did not interfere with mite feeding and feces excretion (Fig. 1B). When mites had the choice to feed on Col-0 or *cyp79b2 cyp79b3* leaves, they showed a strong preference for feeding on *cyp79b2 cyp79b3* leaves (48 and 43 feces associated with feeding on *cyp79b2 cyp79b3* leaves relative to 13 and 23 associated with Col-0 leaves, when *cyp79b2 cyp79b3* leaves were non-stained or blue, respectively), Fig. 1B. Importantly, mites fed on Col-0 leaves even when *cyp79b2 cyp79b3* leaves were present, indicating that Trp-derived metabolites do not deter mites from probing fully-defended leaves. To further characterize the impact of Trp-derived metabolites on mites we performed a no-choice feeding experiment that included mite transfer between leaves of different genotypes. Mites were allowed to feed on leaf 1 for 18 h and were subsequently transferred to leaf 2 for 24 h, upon which the number of feces and eggs was recorded (Fig. 1C). Consistent with the previous report (Zhurov et al., 2014), mites that exclusively fed on *cyp79b2 cyp79b3* leaves deposited about 3 times more feces (measuring mite feeding) and eggs (measuring mite fitness) relative to mites that fed on Col-0 leaves. When mites were transferred from *cyp79b2 cyp79b3* to Col-0, they deposited the similar number of feces and eggs as mites that exclusively fed on Col-0, highlighting the immediate impact of Trp-derivatives on mite feeding. When mites were transferred from Col-0 to *cyp79b2 cyp79b3*, they deposited a slightly but significantly lower number of feces and eggs than mites fed only on *cyp79b2 cyp79b3*, Fig. 1C, demonstrating that the effects of Trp-derived metabolites quickly diminished once they are removed from a diet. The overall similarity between numbers of feces and eggs deposited by mites irrespective of leaf genotypes indicates that Trp-derived metabolites do not impair mite’s ability to acquire nutrients from ingested plant cell content. Rather, our data point to their feeding suppressant effect, causing cessation or slowing of adult female mite feeding.

**Figure 1.**
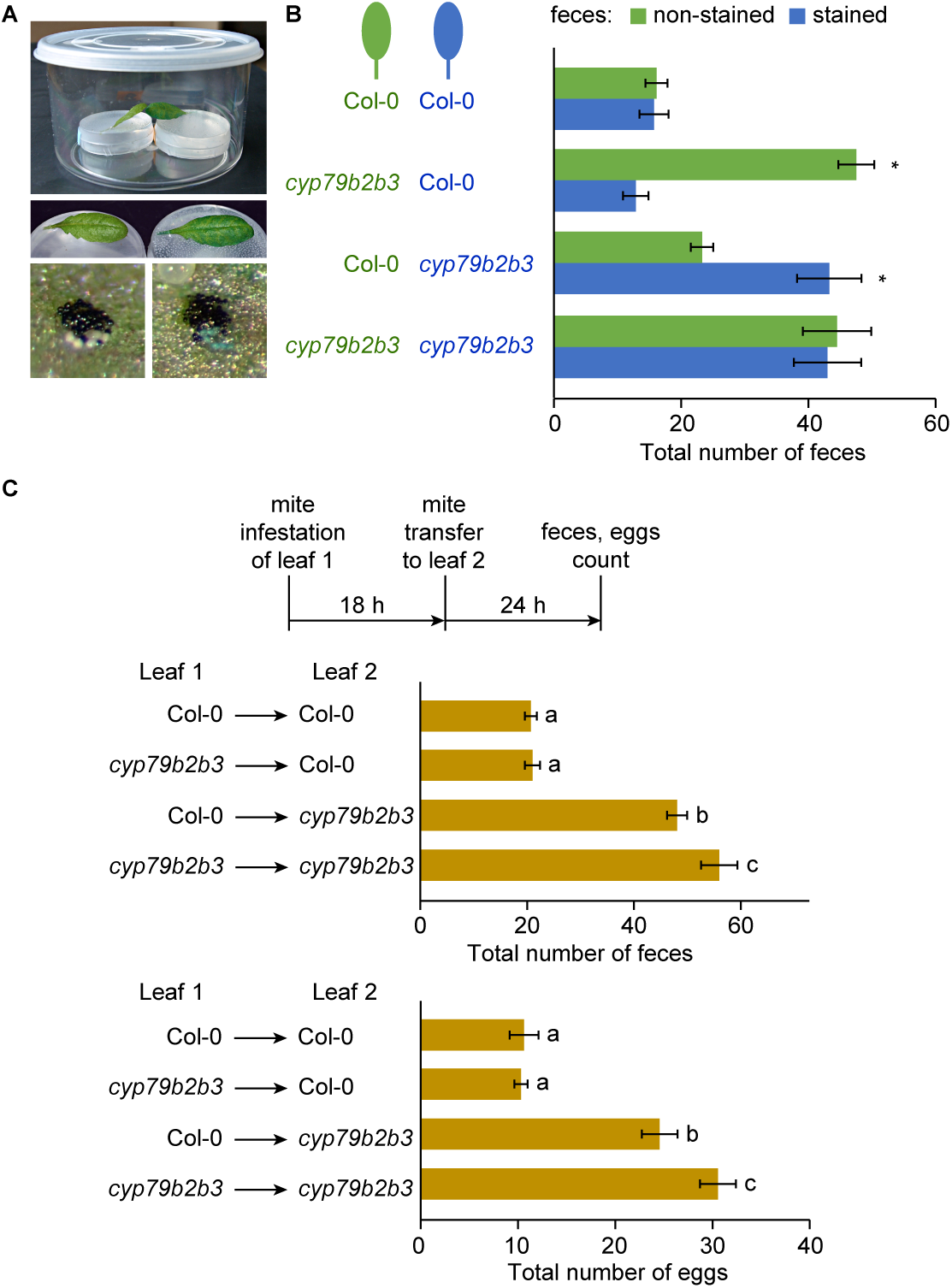
Trp-derived metabolites suppress mite feeding and fecundity. **A**, Experimental set-up for the mite feeding experiment where mites were given the choice to feed on Col-0 or *cyp79b2 cyp79b3* (*cyp79b2b3*) leaves. To track leaves mites fed on, one leaf was supplemented with blue dye and the other remained unstained. Mites feeding on the blue leaf produced blue feces. **B**, The total number of blue and non-stained feces excreted by 10 mites after 48 h. Asterisks indicate a deviation from a 1:1 ratio of non-colored:blue feces (p ≤ 0.05). **C**, The effectiveness of Trp-derived metabolites upon mite transfer between Col-0 and *cyp79b2 cyp79b3* (*cyp79b2b3*) leaves. Ten mites were added to the first leaf and 18 h later, they were transferred to the second leaf. The total number of feces and eggs was scored on the second leaf 24 h after transfer. Significant differences (p ≤ 0.05) are indicated by different letters. (B-C) Experiments were performed in five biological replicates/trial and in three independent trials. Data represent the mean ± SE of three trials.

### I3M is sufficient to restore Arabidopsis defenses against mite infestation in *cyp79b2 cyp79b3* mutant plants

Tryptophan-derived metabolites are synthesized through CYP71A12, CYP71A13, and CYP83B1 pathways (Zhao et al., 2002; Glawischnig 2007; Rajniak et al., 2015; Pastorczyk et al., 2020). To test if metabolites produced through CYP71A12 and/or CYP71A13 pathways affect mite fitness we compared mite fecundity upon feeding on the *cyp71a12* and *cyp71a13* single mutants and *cyp71a12 cyp71a13* (*cyp71a12a13*) double mutant, *cyp79b2 cyp79b3*, and Col-0. As seen in Figure 2A, mite fecundity was similar when they fed on mutant and Col-0 plants, indicating that Trp-derived metabolites synthesized through CYP71A12- and CYP71A13-pathways are not required for the Arabidopsis defense against *T. urticae*. To test if the third, CYP83B1-dependent, indole glucosinolate pathway is sufficient to restore *cyp79b2 cyp79b3* defenses against mites, we infiltrated 2.4 mM or 4.8 mM I3M to *cyp79b2 cyp79b3* and Col-0 detached leaves and subsequently challenged them with *T. urticae*, whose fecundity was determined 48 h later (Fig. 2B and C, and Supplemental Fig. S1). The treatment restored physiological levels of I3M in *cyp79b2 cyp79b3* and increased levels of I3M in Col-0 treated leaves (Fig. 2B). The addition of I3M to *cyp79b2 cyp79b3* leaves fully restored Arabidopsis defenses against mites, Fig. 2C and Supplemental Fig. S1, indicating that I3M is sufficient to establish Trp-derived defenses against *T. urticae*. The supplementation of Col-0 leaves with I3M did not affect mite fecundity, implying that physiological levels of I3M are saturating defenses that could not be further enhanced by a further increase of I3M levels (Fig. 2C).

**Figure 2.**
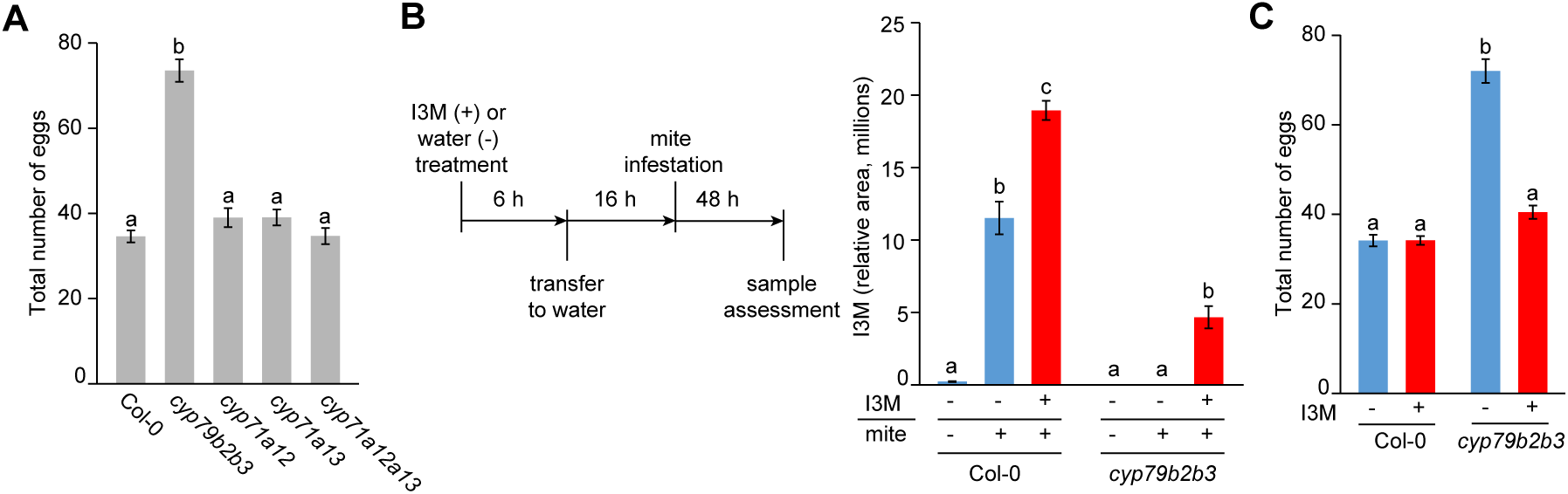
The contribution of individual Trp-derived metabolic pathways to Arabidopsis defenses against mites. **A**, Spider mite fecundity upon feeding on Col-0, *cyp79b2 cyp79b3* (*cyp79b2b3*), *cyp71a12, cyp71a13* and *cyp71a12 cyp71a13* (*cyp71a12a13*) leaves. The total number of eggs per leaf was recorded 48 h after the addition of 10 mites/leaf. **B**, Levels of I3M in Col-0 and *cyp79b2 cyp79b3* (*cyp79b2b3*) leaves supplemented with I3M and infested with 10 mites. (+I3M), leaves supplemented with solution of 2.4 mM I3M for 6 h and kept in water for 16 h before mite addition; (+ mite), leaves challenged with 10 mites for 48h; (-I3M/-mite), untreated leaves immediately frozen after being cut from intact plant. **C**, The effect of I3M supplementation to Col-0 and *cyp79b2 cyp79b3* (*cyp79b2b3*) leaves on total number of eggs laid by 10 mites 48 h after the infestation. A-C, Experiments were performed in at least five biological replicates/trial and in three independent trials. Data represent the mean ± SE of three trials. Significant differences (p ≤ 0.05) are indicated by different letters.

### Multiple indole glucosinolates defend Arabidopsis leaves against mite herbivory

In Col-0 plants, endogenous I3M is oxidized and methylated to give rise to 1MO-I3M and 4MO-I3M. To determine whether exogenously supplied I3M was processed in *cyp79b2 cyp79b3* leaves infested by *T. urticae*, we measured levels of 1MO-I3M and 4MO-I3M in these leaves using an HPLC-MS. As expected, IG metabolites were undetectable in water-treated *cyp79b2 cyp79b3* leaves, regardless of mite infestation status (Fig. 3A). However, in *cyp79b2 cyp79b3* leaves supplemented with I3M, the levels of 1MO-I3M and 4MO-I3M reached about 100 and 50%, respectively, of levels found in mite-infested Col-0 leaves kept in water.Therefore, the I3M infiltrated into *cyp79b2 cyp79b3* leaves was partially processed by CYP81 enzymes and IG methyltransferases into 1MO-I3M and 4MO-I3M, raising the question of which of these metabolites have defensive properties against *T. urticae*.

**Figure 3.**
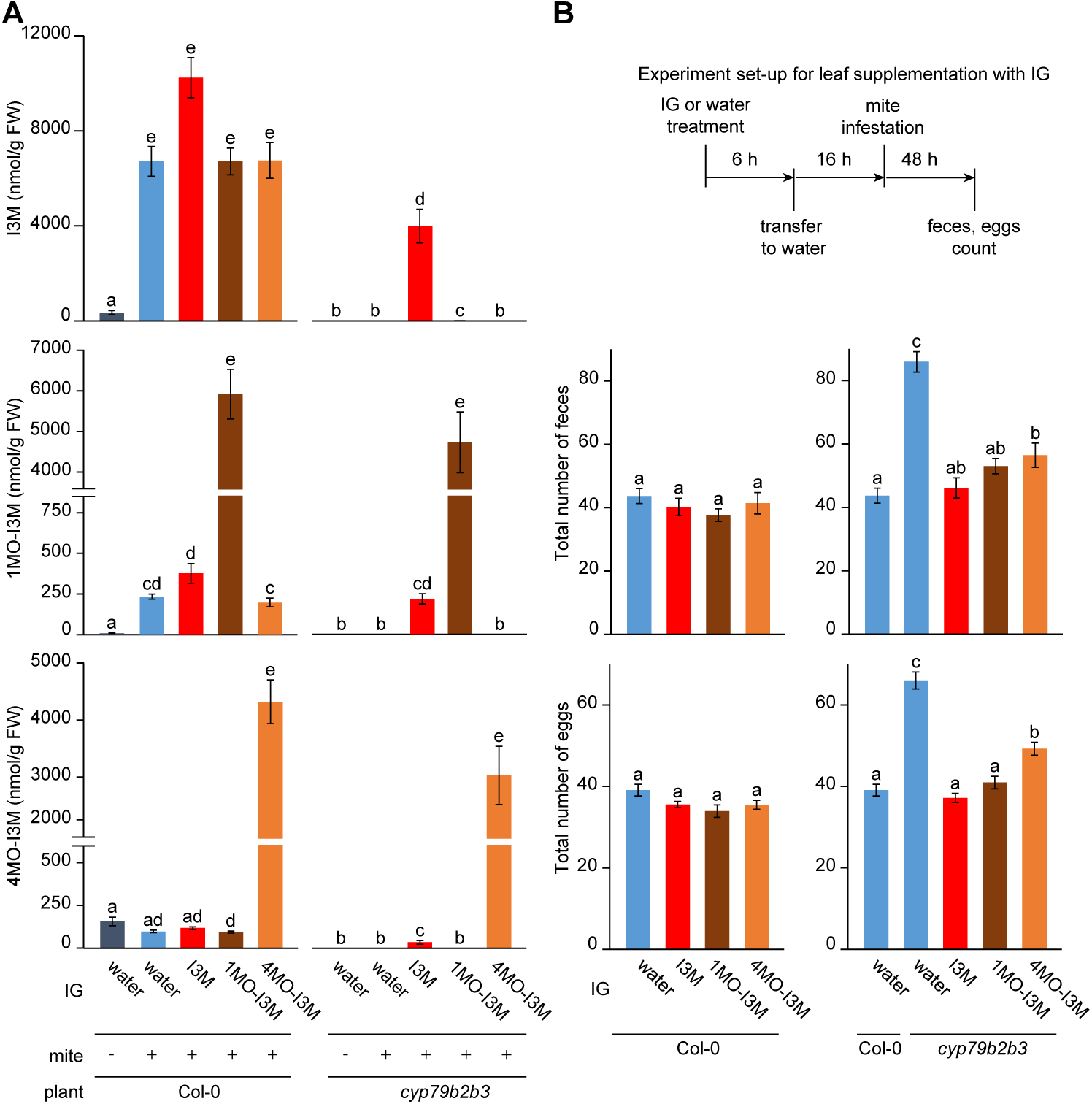
Supplementation with I3M, 1MO-I3M or 4MO-I3M indole glucosinolates fully restores defenses in *cyp79b2 cyp79b3* leaves. **A**, Levels of I3M, 1MO-I3M, and 4MO-I3M in Col-0 and *cyp79b2 cyp79b3* (*cyp79b2b3*) leaves supplemented with IGs. (water/-mite), untreated leaves immediately frozen after being cut from intact plant; (+mite), leaves challenged with mites for 48h. Values were log2 transformed for statistical analysis. **B**, Mite fitness upon feeding on Col-0 and *cyp79b2 cyp79b3* (*cyp79b2b3*) leaves supplemented with I3M, 1MO-I3M or 4MO-I3M. The total numbers of deposited feces (top panels) and eggs (bottom panels) were recorded 48 h after the addition of 10 mites per leaf. Experiments were performed in five biological replicates/trial and in four independent trials. Data represent the mean ± SE of four trials. Significant differences (p ≤ 0.05) are indicated by different letters.

To test the ability of I3M-derived metabolites to reduce mite fitness we infiltrated 2.4 mM solution of 1MO-I3M and 4MO-I3M into *cyp79b2 cyp79b3* and Col-0 detached leaves. Relative to the physiological levels found in Col-0 infested leaves kept in water, the addition of 1MO-I3M and 4MO-I3M resulted in a large excess of these compounds in *cyp79b2 cyp79b3* (21 and 31 fold increase, respectively) and Col-0 leaves (26 and 44 fold change, respectively) (Fig. 3A). Infiltrated leaves were inoculated with ten mites per leaf and the number of eggs and the number of feces were scored two days later, Fig. 3B. The supplementation of Col-0 leaves with I3M, 1MO-I3M, and 4MO-I3M did not affect mite feeding and fecundity, indicating that levels of indole glucosinolates in Col-0 are sufficient to ensure maximal defenses (Fig. 3B). On the contrary, mites deposited significantly fewer feces and eggs on *cyp79b2 cyp79b3* leaves supplemented with either I3M, 1MO-I3M, or 4MO-I3M relative to *cyp79b2 cyp79b3* control leaves kept in water. Of the three indole glucosinolates, supplementation of *cyp79b2 cyp79b3* leaves with I3M and 1MO-I3M reduced the number of feces and eggs to levels seen in Col-0, thus, fully restoring defenses in *cyp79b2 cyp79b3* leaves, Fig. 3B. Mites feeding on 4MO-I3M treated *cyp79b2 cyp79b3* leaves deposited 29 % and 26 % more feces and eggs respectively relative to Col-0. These results demonstrate that I3M and 1MO-I3M, and to a slightly lesser extent 4MO-I3M, can curtail mite feeding and oviposition on Arabidopsis leaves.

### Intact I3M, 1MO-I3M, and 4MO-I3M do not affect mite fitness

I3M, 1MO-I3M and 4MO-I3M glucosinolates require further modifications for anti-herbivory activity. They can be activated by a plant (Barth and Jander 2006) or herbivore gut (Beran et al., 2014) myrosinase, or by the spontaneous breakdown in the gut (Kim and Jander 2007). To discriminate the mode of glucosinolate activation in the *Arabidopsis* - *T. urticae* interaction, we first tested if unmodified glucosinolate compounds affect mite fitness. We fed mites with 0.23, 2.3, or 4.6 mM I3M solutions for 19 h, after which we transferred them to bean leaves for fecundity measurements at 24 and 48 hours, Fig. 4A. Direct delivery of I3M did not affect mite fitness and resulted in similar mite fecundity in treated and control mites (Fig. 4A). To test if continuity of mite exposure to intact indole glucosinolates is required for their effectiveness and to mimic their normal intake through ingestion, we supplemented bean leaf disks with water, I3M, 1MO-I3M, or 4MO-I3M and subsequently examined mite feeding and oviposition over 24 h, Fig. 4B. I3M, 1MO-I3M, or 4MO-I3M extracted from treated bean leaves were stable throughout the experiment, indicating that mites were continuously exposed to constant and high levels of these metabolites, Supplemental Fig. S2. Mites deposited similar numbers of feces and eggs on treated and control bean leaf disks (Fig. 4B), demonstrating that infiltrated indole glucosinolates into bean leaves were ineffective against mites. Overall, these data indicate that I3M, 1MO-I3M, or 4MO-I3M cannot be activated by bean beta-glucosidases or in the mite gut. We therefore considered whether Arabidopsis myrosinases mediate indole glucosinolate activation.

**Figure 4.**
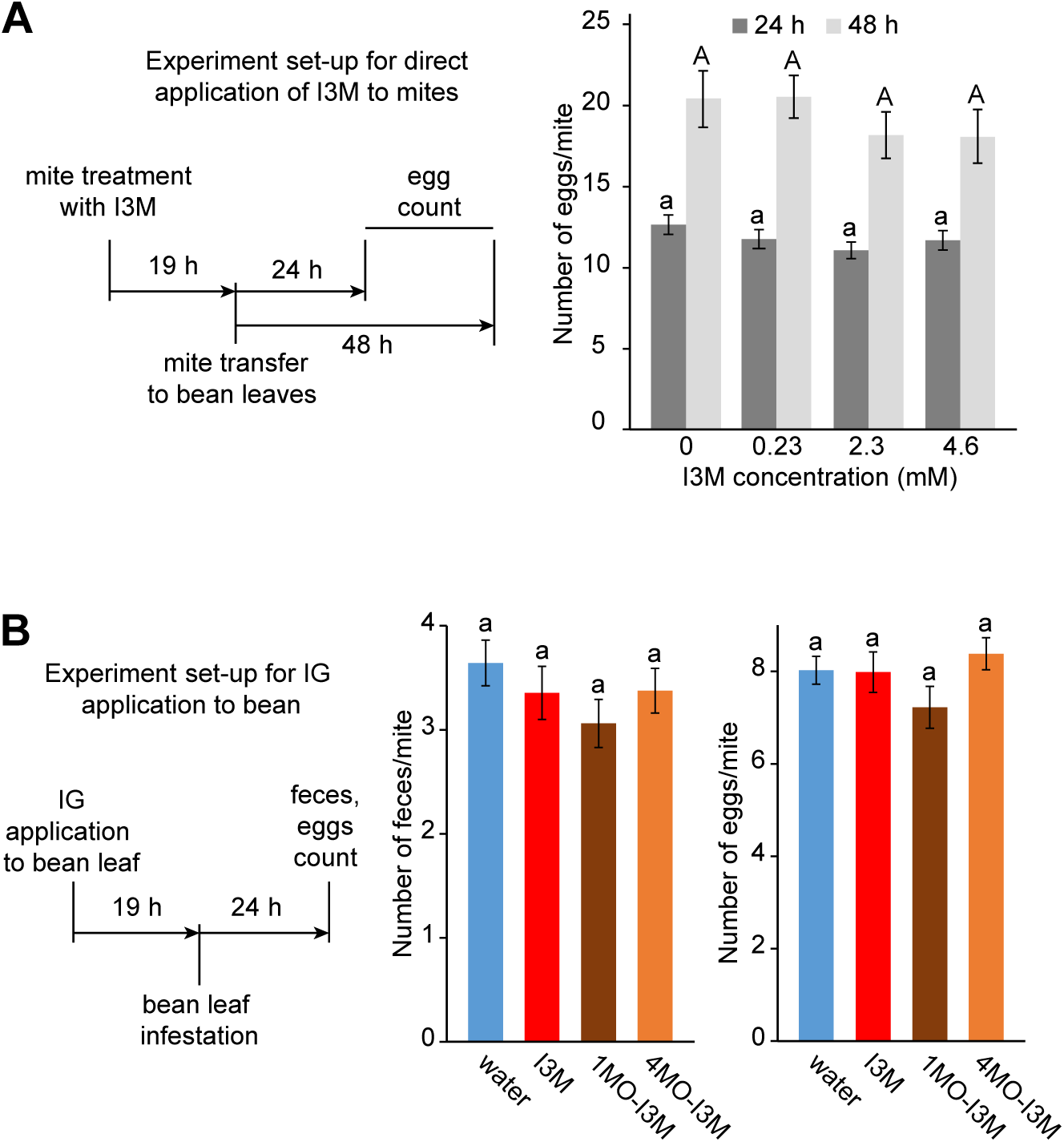
Intact I3M, 1MO-I3M, and 4MO-I3M are not efficient against mites. **A**, Mite fecundity upon direct application of 0.23, 2.3 or 4.6 mM I3M to mites. Mites were treated for 19 h with I3M solutions and were subsequently transferred to bean leaves. Mite fecundity was determined at 24 and 48 h after treatment. **B**, The total number of deposited feces and eggs over 24 h of feeding on bean leaf disk treated with 2.4 mM of I3M, 1MO-I3M, or 4MO-I3M. Experiments were performed in three (in A) and five (in B) biological replicates/trial and in three independent trials. Data represent the mean ± SE of three trials. Significant differences (p ≤ 0.05) are indicated by different letters.

### Classical myrosinases TGG1 and TGG2 are required for indole glucosinolate activity

Classical myrosinases TGG1 and TGG2 are the main Arabidopsis enzymes catalyzing glucosinolate hydrolysis (Barth and Jander 2006). To test the requirement of TGG1 and TGG2 for Arabidopsis defenses against mite herbivory, we compared mite fitness parameters upon their feeding on *tgg1 tgg2* mutant plants relative to Col-0. Mites laid 28% more eggs on *tgg1 tgg2* than on the Col-0 plants (Fig. 5A), caused 89% greater damage (Fig. 5B), required less time to progress through the larval developmental stage (from 5.2 to 4.5 days, Fig. 5C), and had 39% lower larval mortality (Fig. 5D) in the absence of TGG1 and TGG2 activity, firmly supporting the requirement of TGG1 and TGG2 for the establishment of Arabidopsis defenses. If TGG1 and TGG2 were the only factors required for the activation of indole glucosinolates, then the loss of their activity is expected to have the same impact on plant defenses as a loss of indole glucosinolate biosynthesis in *cyp79b2 cyp79b3* plants. However, mites caused 30% greater damage on *cyp79b2 cyp79b3* than on *tgg1 tgg2* mutant plants (Fig. 5B), further accelerated their development from 4.5 to 3.3 days (Fig. 5C), and reduced larval mortality by an additional 25% (Fig. 5D). On *cyp79b2 cyp79b3* relative to Col-0 leaves, mites caused 157% greater damage and larvae mortality was reduced by 82.5%. Therefore, TGG1 and TGG2 contribute to ∼50% of indole glucosinolate activity, suggesting the existence of additional factors that are required for the establishment of indole glucosinolate defenses. We tested the requirement of PEN2, an atypical myrosinase shown to metabolize IGs (Bednarek et al., 2009), however, loss of its function in the *pen2-1* mutant did not affect mite fecundity, Fig. 5E.

**Figure 5.**
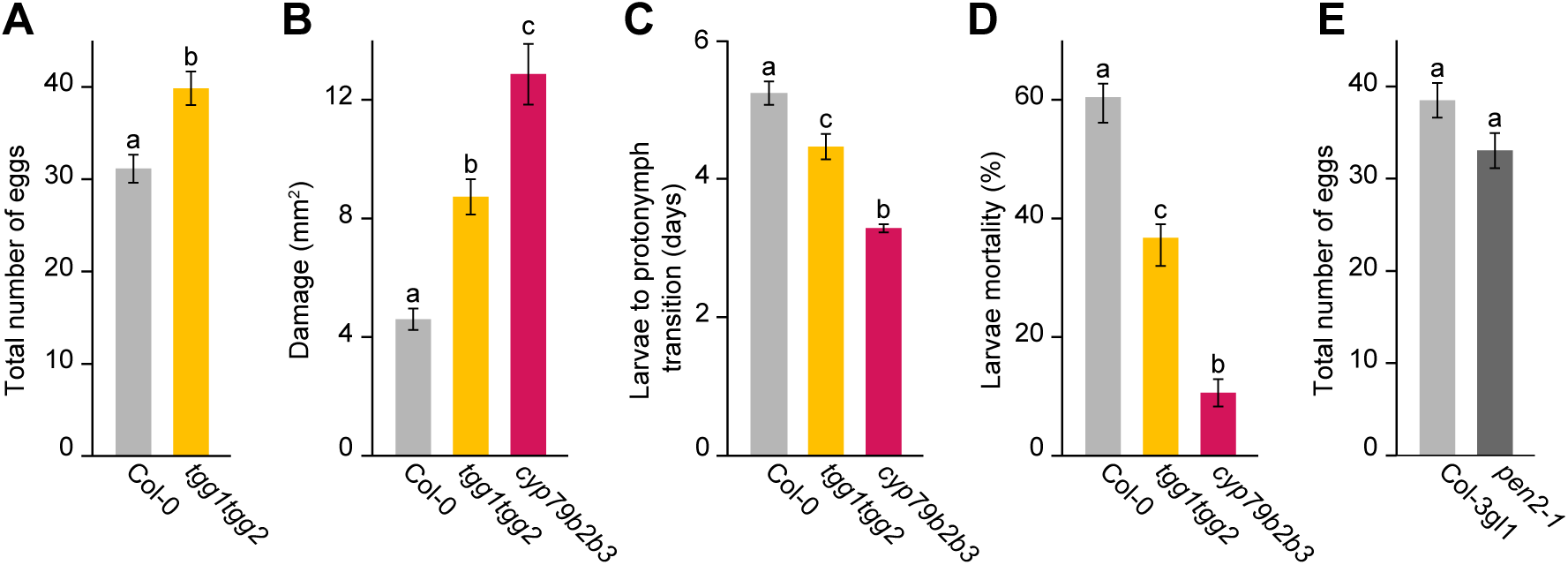
The Arabidopsis myrosinases TGG1 and TGG2 are required for Arabidopsis defense against mites. **A**, Fecundity of 10 mites upon feeding on Col-0 and *tgg1 tgg2* leaves for 48 h. **B-D**, Comparison of mite damage and fitness upon feeding on Col-0, *tgg1 tgg2*, and *cyp79b2 cyp79b3* (*cyp79b2b3*) leaves. **B**, Leaf damage resulting from feeding of 10 mites per plant over three days. **C**, Time required for larvae to become nymphs. **D**, Larval mortality. **E**, Fecundity of 10 mites upon feeding on Col-3 gl1 and *pen2-1* leaves for 48 h. Experiments were performed in five (in A, C-E) and ten (in B) biological replicates/trial and in three independent trials. Data represent the mean ± SE of three trials. Significant differences (p ≤ 0.05) are indicated by different letters.

### The indole glucosinolates are part of the wider MYC2/MYC3/MYC4-regulated defenses against mites

The *myc2 myc3 myc4* mutant plants lack a wide range of JA-regulated Arabidopsis responses including indole glucosinolates (Schweizer et al., 2013). To investigate the relative contribution of indole glucosinolates to JA-regulated defenses against mites, we compared mite fecundity when they fed on *myc2 myc3 myc4* (that lack JA-regulated defenses), *cyp79b2 cyp79b3* (that lack indole glucosinolate defenses), and fully defended Col-0 plants. Mite fecundity was five- and two-fold higher on *myc2 myc3 myc4* and *cyp79b2 cyp79b3*, respectively, relative to Col-0 (Fig. 6A), establishing the existence of a wider array of JA-regulated Arabidopsis defenses against mite herbivory that are mediated through MYC2/MYC3/MYC4 signaling. To directly test the contribution of indole glucosinolates to MYC2/MYC3/MYC4*-*regulated defenses, we supplemented *myc2 myc3 myc4* with I3M. Infiltration of I3M restored I3M and 4MO-I3M to levels measured in mite-infested Col-0 leaves kept in water (Figure 6B). However, 1MO-I3M was undetectable, demonstrating that 1MO-I3M formation upon mite feeding fully depends on MYC2/MYC3/MYC4 transcription factors. Mite oviposition decreased by ∼35% and 42% when fed on *myc2 myc3 myc4* leaves supplemented with I3M or 1MO-I3M compared to *myc2 myc3 myc4* untreated leaves (Fig. 6C and Supplemental Fig. S3), indicating that indole glucosinolates are prominent Arabidopsis defensive compounds against mite feeding. However, I3M and 1MO-I3M only partially complemented defenses in *myc2 myc3 myc4*, confirming the existence of additional, indole glucosinolate-independent, defensive compounds that restrict *T. urticae* herbivory.

**Figure 6.**
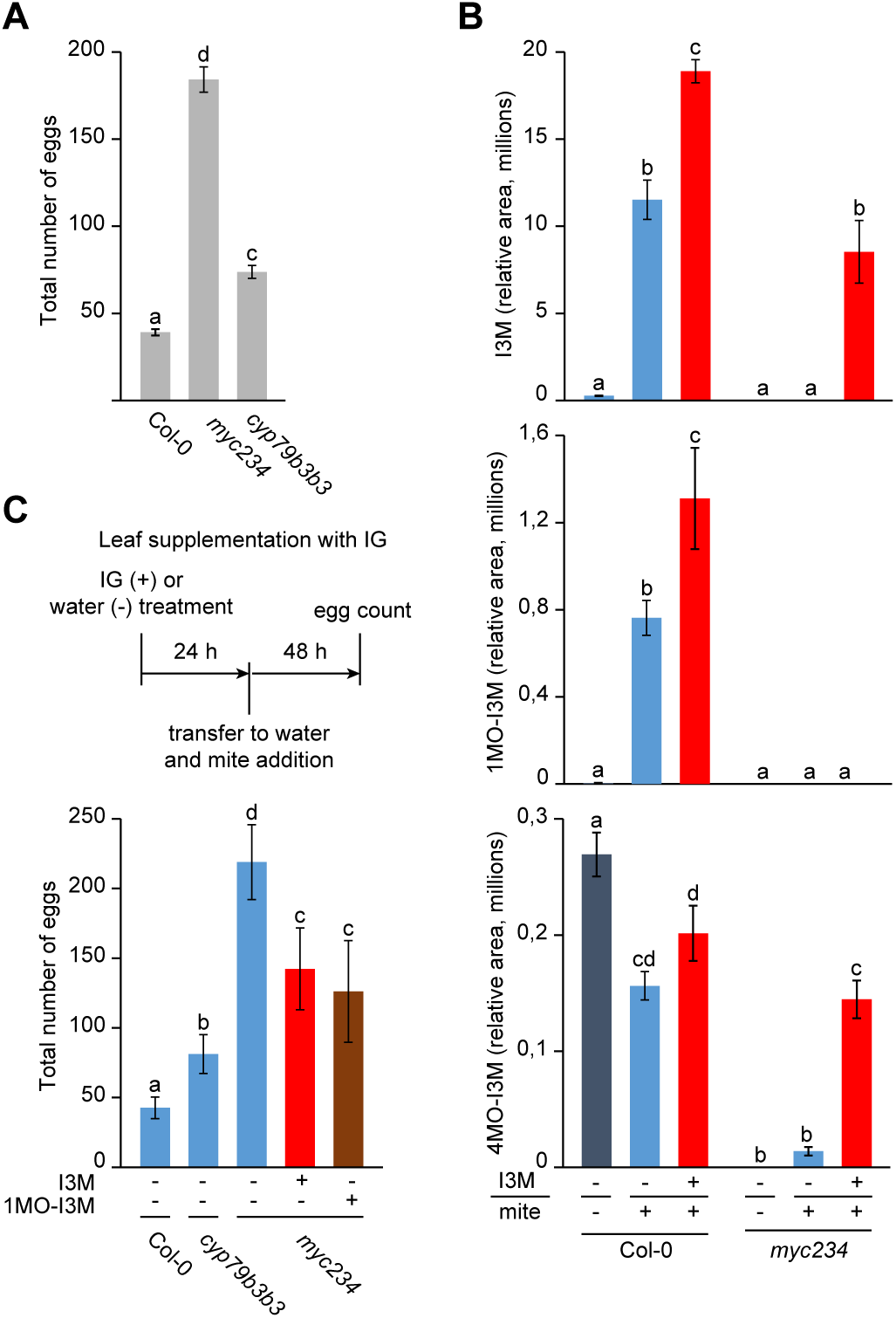
MYC2 MYC3 MYC4 are required for indole glucosinolate-mediated and Trp-independent defenses against mites in Arabidopsis. **A**, Fecundity of 10 mites feeding for 48 h on Col-0, *myc2 myc3 myc4* (*myc234*) and *cyp79b2 cyp79b3* (*cyp79b2b3*) leaves. **B**, Levels of I3M, 1MO-I3M and 4MO-I3M in Col-0 and *myc2 myc3 myc4* (*myc234*) leaves supplemented with 2.4 mM I3M for 6 h and kept in water for 16 h before mite addition. (-I3M/-mite), untreated leaves were immediately frozen after being cut from intact plant; (+ mite) leaves challenged with mites for 48h. **C**, Fecundity of 10 mites upon feeding for 48 h on Col-0, *cyp79b2 cyp79b3* (*cyp79b2b3*) and *myc2 myc3 myc4* (*myc234*) leaves supplemented with 4.8 mM I3M or 1MO-I3M for 24 h before mite addition. Experiments were performed in five biological replicates/trial and in three independent trials. Data represent the mean ± SE of three trials. Significant differences (p ≤ 0.05) are indicated by different letters.

## DISCUSSION

Camalexin, cyanogenic 4-OH-ICN metabolite, and indole glucosinolates are three classes of known tryptophan-derived defense metabolites. Among them, camalexin and 4-OH-ICN, produced through CYP71A12 and CYP71A13, were shown to act as specialized defense compounds against aphids and a variety of plant pathogens (Thomma et al., 1999; Bohman et al., 2004; Ferrari et al., 2007; Sanchez-Vallet et al., 2010; Schlaeppi et al., 2010; Kettles et al., 2013; Glindemann et al., 2019; Pastorczyk et al., 2020). Here, we demonstrated that defense compounds synthesized through CYP71A12- and CYP71A13-dependent pathways are not protecting Arabidopsis from mite herbivory, Fig. 2A, even though levels of camalexin increase upon mite feeding (Zhurov et al., 2014). Instead, we showed that I3M, a parental metabolite of the indole glucosinolate pathway, can fully restore Trp-derived defenses controlling mite herbivory in otherwise defenseless *cyp79b2 cyp79b3* plants, Fig. 2C. We further established that exogenously supplied I3M is converted into 1MO-I3M and 4MO-I3M indole glucosinolates in leaf tissues, and that all three metabolites act redundantly to control mite infestation, Fig. 3.

Upon mite feeding, both endogenous and exogenously supplied I3M are preferentially converted to 1MO-I3M rather than to 4MO-I3M, Fig. 3. This favors Arabidopsis defenses against mites, as 1MO-I3M had greater suppressant effects on mites than 4MO-I3M (Fig. 3A). In contrast, in response to *Myzus persicae* feeding, I3M is exclusively modified to produce aphid-deterrent 4MO-I3M (Kim and Jander 2007). The control of metabolic fluxes within the indole glucosinolate biosynthetic pathway is achieved through transcriptional regulation of specific CYP81F enzymes and IG methyltransferases (IGMTs) (Winter et al., 2007; Pfalz et al., 2016). Conversion of I3M into 1MO-I3M and 4MO-I3M is carried out by complementary CYP81F and IGMT enzymes (Pfalz et al., 2016). Consistent with the dependence of 1MO-I3M synthesis on the MYC2/MYC3/MYC4, Fig. 6, the expression of *CYP81F4* and *IGMT5* that are required for the conversion of I3M into 1MO-I3M is JA-dependent (Winter et al., 2007; Schweizer et al., 2013; Sun et al., 2013). As mite feeding induces the accumulation of JA, whose effects are mediated by MYC2/MYC3/MYC4, it follows that mites also induce the synthesis of 1MO-I3M. In contrast, in response to *M. persicae* herbivory, Arabidopsis plants do not accumulate JA and the expression of indole glucosinolate biosynthetic genes is suppressed (De Vos et al., 2005; Giordanengo et al., 2010; Sun et al., 2013; Appel et al., 2014; Foyer et al., 2015). Consequently, aphid feeding does not trigger the accumulation of 1MO-I3M.

During vegetative development, Arabidopsis plants synthesize indole glucosinolates in both root and leaf tissues and can bidirectionally transport them between these organs (Andersen et al., 2013). Upregulation of indole glucosinolate biosynthetic genes in leaves in response to mite feeding indicates that at least a portion of defensive glucosinolates is synthesized in leaves (Zhurov et al., 2014). The vasculature has been proposed to facilitate indole glucosinolate long-distance transport, even though transporters that enable movement of indole glucosinolates between vasculature and the apoplast are currently unknown (Andersen et al., 2013). The complete restoration of defenses in *cyp79b2 cyp79b3* leaves that were being supplemented with I3M through the petiole, expected to be taken up through the xylem, Fig. 2 and 3, demonstrates that leaf-imported I3M can defend leaf tissues against mites. Importantly, the infiltrated I3M must have been taken-up by cells, as it was converted into 1MO-I3M and 4MO-I3M by intracellular CYP81F and IGMT enzymes, Fig. 3. Endogenous I3M is synthesized in leaves by CYP83B1 localized in specialized cells that are adjacent to the phloem (Nintemann et al., 2018) and is stored in vasculature-associated S-cells (Koroleva et al., 2010). It is further distributed throughout the leaf blade with higher accumulation in the abaxial epidermal cells (Madsen et al., 2014). Whether this pattern of indole glucosinolate accumulation changes in response to mite feeding is currently unknown, however, its broad distribution is likely preserved.

Intact glucosinolates are not toxic and require modifications for their defensive activity. TGG1 and TGG2 are major Arabidopsis myrosinases that cleave thioglucoside bonds within the glucosinolate molecules, releasing highly reactive aglucones that are further modified to yield defensive compounds (Rask et al., 2000; Wittstock and Halkier 2002; Barth and Jander 2006; Kissen et al., 2009). High levels of TGG1 and TGG2 accumulate in guard cells in epidermal (Zhao et al., 2008), and in myrosin cells that are localized in the vicinity but in non-overlapping cells relative to glucosinolate synthesizing and storing cells, in phloem parenchyma (Husebye et al., 2002; Kissen et al., 2009; Li and Sack 2014; Shirakawa et al., 2014; Burow and Halkier 2017). The physical separation between glucosinolate- and myrosinase-storing cells led to a hypothesis that tissue maceration is required to enable their interaction and the generation of glucosinolate breakdown products. Consistent with this model and the requirement of TGG1 and TGG2 for the activation of aliphatic glucosinolates, Arabidopsis defenses against generalist chewing herbivores like *Manduca sexta* and *Trichoplusia ni* (Müller et al., 2010), are dependent on the activity of TGG1 and TGG2 (Barth and Jander 2006). However, the fitness of the Hemiptera *Myzus persicae* and *Brevicoryne brassicae* that are sensitive to indole glucosinolates is not affected by these myrosinases (Barth and Jander 2006). It has been postulated that aphid feeding, involving intercellular movement of the stylet before it reaches the phloem sieve elements (Tjallingii and Hogen Esch 1993), avoids rupturing of the myrosinase containing cells and thus prevents contact between TGG1 and TGG2 and glucosinolates (Kim et al., 2008). However, IGs are broken down post-ingestion in the aphid gut where they form conjugates that restrict aphid herbivory (Kim and Jander 2007). Whether the aphid gut contains enzymes with myrosinase activity, like *Phyllotreta striolata* (flea beetle) (Beran et al., 2014), or indole glucosinolates undergo spontaneous breakdown is at present unknown. Similar to aphids, mites use stylets to feed from individual mesophyll parenchyma cells (Bensoussan et al., 2016). Intact indole glucosinolates did not affect mite fitness (Fig. 4), indicating that the *T. urticae* gut lacks myrosinase activity and does not destabilize these metabolites. Instead, we found that the Arabidopsis myrosinases TGG1 and TGG2 are required to limit mite proliferation (Fig. 5). This is surprising, as mites, like aphids, are not expected to feed from the myrosinase-containing cells in phloem parenchyma. Whether mites sample some of the cellular content of guard cells as they sometimes protrude their stylets through a stomatal opening, and thus ingest some of the TGGs, is not known. In that case, the well-known “mustard oil bomb” system could be reconstituted in the mite gut. Alternatively, TGG myrosinases could be expressed in mesophyll parenchyma cells at a low level and thus may have evaded detection by *in situs*, promoter fusions, and antibody stainings (Xue et al., 1995; Husebye et al., 2002; Barth and Jander 2006; Kissen et al., 2009; Shirakawa and Hara-Nishimura 2018). This scenario could enable the activation of the “mustard oil bomb” within a single cell that mites consume. Alternatively, the effect of TGG1 and TGG2 may be indirect through the modification of the morphological and chemical properties of pavement and stomatal cells (Ahuja et al., 2016) that may hinder stylet penetration through the epidermis.

Mite fitness was greater on *cyp79b2 cyp79b3* than on *tgg1 tgg2* mutant plants (Fig. 5), which indicates the involvement of additional Arabidopsis factors in the generation of indole glucosinolate-dependent defensive compounds. For example, several other enzymes were shown to process indole glucosinolates (Bednarek et al., 2009; Clay et al., 2009; Nakano et al., 2017). One of them, PEN2 is required for Arabidopsis defenses against several pathogens (Lipka et al., 2005; Bednarek et al., 2009; Clay et al., 2009), but is dispensable for restricting mite fitness, Fig. 5E. PYK10 and BGLU18 are additional beta-glucosidases capable of hydrolyzing I3M and 4MO-I3M (Nakazaki et al., 2019; Sugiyama and Hirai 2019). Whether they are required for the Arabidopsis defenses against mites is currently not known. Regardless, Arabidopsis factors required for the generation of indole glucosinolate-dependent defensive compounds that act downstream of IG biosynthesis appear to be limiting plant defense against mites, as the excess of I3M, 1MO-I3M and 4MO-I3M in IG-supplemented Col-0 leaves did not increase Arabidopsis defenses against mites (Fig. 2 and 3). The processing of I3M, 1MO-I3M, and 4MO-I3M is expected to yield multiple active compounds that may affect mites directly or may induce the production of other defense compounds (Bednarek et al., 2009; Clay et al., 2009; Matern et al., 2019). In addition to IG-derived defense compounds, Arabidopsis plants likely have additional phytochemicals capable of restricting mite herbivory. They are dependent on MYC2/MYC3/MYC4 but are synthesized independently of CYP79B2/CYP79B3, Fig. 6. Thus, our results demonstrate that Arabidopsis plants trigger at least two independent defense pathways that result in a complex and diverse array of chemical defenses against *T. urticae* herbivory.

### Conclusions

The *cyp79b2 cyp79b3* mutant plants lack Trp-derived metabolites and are more sensitive to *T. urticae*. Here, we demonstrated that the Trp-derived metabolites synthesized through the indole glucosinolate pathway are efficient against mites. Three indole glucosinolates, I3M, 1MO-I3M, and 4MO-I3M, are able to complement the *cyp79b2 cyp79b3* mutant leaves and restore feeding suppressant defenses against mites. Intact indole glucosinolates are ineffective against mites. They require TGG1, TGG2 and other, at present unknown, myrosinases to restrain mite proliferation. Indole glucosinolates are part of the wider MYC2/MYC3/MYC4-regulated defenses against mites, indicating the complexity of Arabidopsis defenses against *T. urticae* herbivory.

## MATERIALS AND METHODS

### Plant materials and growth conditions

The *Arabidopsis thaliana* wild-type seeds were obtained from the *Arabidopsis Biological Resource Center* for Columbia-0 (Col-0) and H. Ghareeb (Göttingen University) for Col-3 gl1. The seeds of *myc2 myc3 myc4* were kindly provided by R. Solano (Universidad Autónoma de Madrid), *cyp79b2 cyp79b3* by B. A. Halkier (University of Copenhagen), *cyp71a12, cyp71a13* and *cyp71a12 cyp71a13* by E. Glawischnig (Technical University of Munich), *pen2-1* by H. Ghareeb (Göttingen University), and *tgg1 tgg2* by G. Jander (Cornell University). All Arabidopsis mutants are in the Col-0 background, except *pen2-1* which is in the Col-3 gl1 background. Plants were grown at 21-22°C, with 50% relative humidity and a short-day (10 h day/14 h night) photoperiod. All Arabidopsis plants used for experiments were 4 to 5-week-old. Bean plants (*Phaseolus vulgaris* cultivar ‘California Red Kidney’; Stokes, Thorold, ON) were grown at 24°C, 55% relative humidity, and with a long-day (16 h day/8 h night) photoperiod. All bean plants used for experiments as well as the maintenance of the spider mite population were 2-week-old.

### Spider mite strain and rearing conditions

The London reference *Tetranychus urticae* strain was reared on bean plants at 24°C, 55% relative humidity, and long-day (16 h day/8 h night) photoperiod, as described previously (Suzuki et al., 2017).

### Mite fecundity tests on detached *Arabidopsis* leaves

The petiole of fully developed leaves was cut and submerged in 10 mL of water contained in a small Petri plate covered with parafilm. Six hours later, each leaf was infested with 10 adult female mites and the Petri plate was covered with a vented lid. The total number of eggs deposited by 10 mites was recorded 48 h following mite infestation. All experiments were performed with at least five leaf replicates per genotype, which were repeated in three independent trials with independent sets of plants.

### Choice experiment

Fully-elongated adult leaves of Col-0 and *cyp79b2 cyp79b3* were cut and each petiole was inserted in a PCR tube containing water or 6% of blue food dye (erioglaucine; McCormick, Sparks Glencoe, MD). After 6 h, leaves were transferred and kept overnight in a small Petri plate set-up described above. One blue-stained and one unstained leaf were then placed into a vented box. 10 adult female spider mites were added to the box. The number of blue feces and unstained feces was recorded 48 h after mite addition. Five biological replicates/trial were performed in three independent trials, with independent sets of plants.

### No-choice experiment

Fully-elongated adult leaves were cut and petioles were inserted in a small Petri plate that contained 10 mL of water and was covered with parafilm. Mites were retained within the Petri dish with a vented lid. Twenty four hours later, leaves (labeled as “leaf 1”) were infested with 10 adult female mites. Eighteen hours following mite application, mites were transferred to a new leaf (“leaf 2”). The numbers of eggs and feces were counted 24 h following mite transfer to leaf 2. Experiments were performed in five biological replicates/trial and three independent trials, with independent sets of plants.

### Fecundity assay on Arabidopsis leaves supplemented with IGs

Fully developed leaves from five-week-old Col-0, *cyp79b2 cyp79b3* or *myc2 myc3 myc4* plants were detached and their petiole was inserted into a solution of I3M, 1MO-I3M, or 4MO-I3M, or water as a control. The 3-indolylmethyl glucosinolate or glucobrassicin (I3M), the neoglucobrassicin (1MO-I3M), and the 4-methoxyglucobrassicin (4MO-I3M) potassium salts were purchased from Extrasynthese (France), with the catalog numbers 2525, 2519 S, and 2522 S, respectively. Depending on the experiment, leaves were treated for either 6 h with 2.4 mM glucosinolate solution and then kept in water overnight before mite infestation, or for 24 h with 4.8 mM glucosinolate solution and were then transferred into the water for immediate mite infestation. After treatment, each leaf (inserted in a small Petri plate covered with parafilm and containing 10 mL of water) was infested with ten adult female mites, which were retained within the plate with a vented lid. The number of eggs and feces was counted 48 h following mite leaf infestation. For each condition of compound supplementation, five biological replicates/trial were performed in three to four independent trials, with independent sets of plants.

### Fecundity assay on bean leaf disks supplemented with IGs

We followed protocol described by Ghazy et al. to deliver IGs to bean leaves (Ghazy et al., 2020). Briefly, bean leaf disks, 1.2 cm in diameter, were excised with a hole puncher from the first pair of leaves of 2-week-old bean plants. A 9.6 µL volume of I3M, 1MO-I3M, or 4MO-I3M solution at 4.6 mM, or of water as a control, was spread on the adaxial side of each disk. The disks were immediately covered with parafilm to avoid leaf desiccation and 3 h later, 10 adult female mites were placed on the adaxial side of each disk, surrounded by wet Kimwipe strips (Kimberly-Clark Professional Kimtech Science Kimwipes) to prevent mite escape. The number of eggs and feces was recorded 24 h after mite addition to the disks. For each compound, five leaf disks were used in each of three independent trials, with independent sets of plants.

### Direct application of IGs to mites

The 3-indolylmethyl glucosinolate (I3M) was applied directly to mites following protocol described in Suzuki et al., 2017. Briefly, a piece of 5 x 5 mm Kimwipes (Kimberly-Clark Professional Kimtech Science Kimwipes) was soaked with 10 µL of I3M solution applied at 0.23 mM, 2.3 mM, or 4.6 mM, or with 10 µL water as a control, and was kept in a sealed Petri plate for 19 h at 20°C. Subsequently, mites were transferred to bean leaf squares laid on a wet filter paper. Fecundity was assessed 24 h and 48 h after mite transfer. One biological replicate comprised one bean leaf square with 10 mites. For each concentration, three biological replicates/trial were performed in three independent trials, with independent sets of plants.

This method of delivery was validated chemically with a coumarin solution (10 mM) and its control solution (10% methanol), Supplemental Fig. S4. Following coumarin treatment, mite mortality was assessed 2 h after transfer on a bean leaf. One biological replicate comprised one bean leaf with ∼30 mites. Three biological replicates/trial were performed in three independent trials, with independent sets of plants.

### Plant damage assay

Ten adult female mites were placed on Col-0, *tgg1 tgg2* and *cyp79b2 cyp79b3* plants. Three days later, the entire rosettes were cut from the roots and scanned using a Canon CanoScan 8600F model scanner at a resolution of 1200 dpi and a brightness setting of +25. Pictures were saved as .jpg files and damage quantification was subsequently performed with Adobe Photoshop 5 (Adobe Systems, San Jose, CA) as described previously (Cazaux et al., 2014). Ten plants per genotype were used per trial. The experiment was performed in three independent trials with independent sets of plants.

### Mortality and larval developmental assays

Fully-elongated adult leaves of Col-0, *tgg1 tgg2* and *cyp79b2 cyp79b3* were cut and each petiole was inserted in a small Petri plate containing 10 mL of water covered with parafilm. Subsequently, 25 newly molted larvae were placed on each leaf and a vented lid was fixed to prevent mite escape. On each day following infestation, the number of larvae, their viability and molting were recorded. The average number of days required for 25 larvae to develop into protonymphs on each detached leaf was used as the data point. Leaves were replaced every other day (day 0, 2, 4, etc.) until all larvae either molted into protonymphs or died. Protonymphs were removed when counted. Five leaves per genotype were infested per trial and the experiment was repeated in three independent trials.

### Indole glucosinolate analysis by HPLC-MS

Metabolites were extracted from frozen leaves in a methanol 70% buffer containing the allylglucosinolate sinigrin (80 µg/mL, (−)-Sinigrin hydrate, Sigma-Aldrich) as an internal standard, with a fresh weight/buffer volume ratio of 100 mg/mL. After grinding the biological sample in the buffer manually with a pestle, metabolites were further extracted by vortexing (1 min) and then by sonication (10 min). Debris was removed by two successive centrifugations at 16160 *g* and the supernatant was analyzed by high-performance liquid chromatography coupled with a time-of-flight mass spectrometry (LC/ TOF MS) using an Agilent 1260 Infinity LC system coupled to an Agilent 6230 TOF system. The Zorbax Eclipse Plus C-18 column Rapid Resolution HT (3 X 100 mm, 1.8 μm, 600 bar, Agilent, USA) was kept at 25°C and the elution was performed with acetonitrile (Optima, Fisher chemical, UK) and water containing formic acid (Sigma-Aldrich, Germany). A gradient of solvent A (H_2_O containing 0.1% formic acid) and solvent B (CH_3_CN 90% in H_2_O, containing 0.1% formic acid) was applied as follows. The initial condition was 5% B in A, which was held for 2 minutes, with the first minute of eluent sent to waste. Metabolites were eluted with a linear gradient to 100% B over 20 minutes. After a 5 minute wash at 100% B, initial conditions of 5% B in A were established over 1 minute followed by a 4 min post run at initial conditions before the next injection. The injection volume was 10 µL for each sample. The flow rate was set to 0.4 mL/min and infused into an Agilent 6230 TOF MS through a Dual Spray ESI source with a gas temperature of 325°C flowing at 8 L/min, and a nebulizer pressure of 35 psi. The fragmentor voltage was set to 175 V with a capillary voltage of 3500 V and a skimmer voltage of 65 V. The instrument was set in negative ESI mode. The negative-ion full-scan mass spectra were recorded over a 85 to 1200 m/z range. The MassHunter Workstation Software Qualitative analysis Version B.05.00 (Agilent Technologies, Inc. 2011) was used for visualizing the chromatograms and peak integration. The compounds of interest were detected by the following ions [M-H]^-^ (theoretical mass, actual mass found) at specific retention times (RT): sinigrin, m/z 358.0272, 358.0314 (RT 1.8 min); I3M, m/z 447.0537, 447.0547 (RT 6.8 min); 4MO-I3M m/z 477.0643, 477.0675 (RT 7.8 min) and 1MO-I3M, m/z 477.0643, 477.0663 (RT 8.8 min). The relative quantification of each metabolite was obtained by correcting the peak area with that of the recovery of the internal standard sinigrin, and is expressed in area units (a.u.). The absolute amount of each IG in plant extracts was further calculated based on standard ranges of synthetic I3M, 4MO-I3M, and 1MO-I3M and the tissue weight, and expressed in nmol/g of fresh weight (F.W.).

### Statistical analysis

Statistical analysis was performed using R software (R Core Team, 2014). For the fecundity tests on Arabidopsis mutant leaves, the no-choice experiment, the fecundity assay on bean leaves supplemented with I3M, the direct application of IGs to mites, and the metabolic analysis of IG solutions used for experiments, we used a two-way ANOVA testing for the main effects of trial, genotype, and any interaction. Interaction terms including trial were included in all statistical analyses to test for reproducibility between trials (Brady et al., 2015). A Tukey’s honestly significant difference (HSD) test was performed following the ANOVA to determine differences between genotypes or between treatments. For the fecundity assays on Arabidopsis leaves supplemented with IGs, count data were analyzed by two-way ANOVA with the interaction of plant genotype and supplemented compound used as the first explanatory variable and trial as the second. No significant effect of the experimental trial was detected, and ANOVA was followed by Tukey’s HSD test. For the two-choice experiment with Col-0 and *cyp79b2 cyp79b3* leaves, after establishing homogeneity of response, count data from individual trials were pooled for the final analysis using goodness-of-fit G-test using R package DescTools (R Core Team, 2014) to assess the statistical significance of feces color deviation from a 1:1 ratio of non-colored:blue feces. For the metabolic analysis of leaves from Arabidopsis and bean supplemented with IGs, and the fecundity assays on Col-0, *cyp79b2 cyp79b3*, and *myc2 myc3 myc4* leaves supplemented with I3M, a three-way ANOVA was performed testing for the main effects of trial, genotype, and treatment. A lack of significant interactions (2 and 3-way) with the other main effects of genotype and treatment signified the data could be combined across trials. The linear model was then simplified by excluding non-relevant, non-significant interactions leaving the 3 main effects and the biologically relevant interaction term between genotype and treatment only. Another three-way ANOVA was performed using the simplified linear model. Differences between genotypes and treatment were determined with a Tukey’s HSD test. The mortality experiment involving coumarin was analysed using a two-way ANOVA testing for the main effects of treatment and trial and any interaction between the two variables. Post-hoc analysis was not required as there were only two treatments to be compared. For the developmental and mortality assays using the *tgg1 tgg2* Arabidopsis mutant, two-way ANOVAs testing for the main effects of trial, genotype, and any interaction between the two variables was used. A Tukey’s honestly significant difference (HSD) test was performed following the ANOVA to determine differences in pairwise comparisons between genotypes.

## Acknowledgments

The authors thank an undergraduate student Emma Somerville for her help with experiments.

## Figure legends

**Supplemental Figure S1. Mite fecundity upon feeding on *cyp79b2 cyp79b3* leaves supplemented with 4**.**8 mM I3M over 24 h**. The total number of eggs per leaf was recorded 48 h after the addition of 10 mites/leaf. Experiment was performed in five biological replicates/trial and in three independent trials. Data represent the mean ± SE of three trials. Significant differences (p ≤ 0.05) are indicated by different letters.

**Supplemental Figure S2. Stability of I3M, 1MO-I3M and 4MO-I3M in supplemented bean leaf disks**. T0, leaf disks collected immediately after application of I3M, 1MO-I3M or 4MO-I3M; T51, leaf disks collected at the end of the experiment, 48 h after mite infestation (51 h after IG application). Experiment was performed in three biological replicates/trial and in three independent trials. Data represent the mean ± SE of three trials. Significant differences (p ≤ 0.05) are indicated by different letters.

**Supplemental Figure S3. I3M supplementation partially rescues defenses in *myc2 myc3 myc4* leaves**. Mite fecundity upon feeding on Col-0, *cyp79b2 cyp79b3* (*cyp79b2b3*) and *myc2 myc3 myc4* (*myc234*) leaves supplemented with 2.4 mM I3M for 6 h and kept in water for 16 h before mite addition. The total number of eggs per leaf was recorded 48 h after the addition of 10 mites/leaf. Experiment was performed in at least five biological replicates/trial and in three independent trials. Data represent the mean ± SE of three trials. Significant differences (p ≤ 0.05) are indicated by different letters.

**Supplemental Figure S4. Mite mortality after direct application of 10 mM coumarin solution to mites**. Experiment was performed in three biological replicates/trial and in three independent trials. Data represent the mean ± SE of three trials. Significant difference (p ≤ 0.05) is indicated by an asterisk.

